# Disease-linked mutations trigger exposure of a protein quality control degron in the DHFR protein

**DOI:** 10.1101/2021.11.04.467226

**Authors:** Caroline Kampmeyer, Sven Larsen-Ledet, Morten Rose Wagnkilde, Mathias Michelsen, Henriette K. M. Iversen, Sofie V. Nielsen, Søren Lindemose, Alberto Caregnato, Tommer Ravid, Amelie Stein, Kaare Teilum, Kresten Lindorff-Larsen, Rasmus Hartmann-Petersen

## Abstract

Degrons are short stretches of amino acids or structural motifs that are embedded in proteins. They mediate recognition by E3 ubiquitin-protein ligases and thus confer protein degradation via the ubiquitin-proteasome system. Well-described degrons include the N-degrons, destruction boxes, and the PIP degrons, which mediate the controlled degradation of various proteins including signaling components and cell cycle regulators. In comparison, the so-called protein quality control (PQC) degrons that mediate the degradation of structurally destabilized or misfolded proteins are not well described. Here, we show that disease-linked DHFR missense variants are structurally destabilized and chaperone-dependent proteasome targets. We systematically mapped regions within DHFR to assess those that act as cytosolic PQC degrons in yeast cells. Two regions, DHFR-Deg13-36 (here Deg1) and DHFR-Deg61-84 (here Deg2), act as degrons and conferred degradation to unrelated fusion partners. The proteasomal turnover of Deg2 was dependent on the molecular chaperone Hsp70. Structural analyses by NMR and hydrogen/deuterium exchange revealed that Deg2 is buried in wild-type DHFR, but becomes transiently exposed in the disease-linked missense variants.

## Introduction

Since evolution has not selected for overly stable proteins, most proteins are only marginally stable under physiological conditions (Bartlett and Radford, 2009; Kim et al., 2013; Maxwell et al., 2005). Consequently, as a result of mutations or stress conditions, proteins are prone to become structurally destabilized or misfolded (Stein et al., 2019). Since such proteins may interact non-specifically with other cell components and be toxic, all cells are equipped with an effective protein quality control (PQC) system that upholds proteostasis by catalyzing the refolding or degradation of aberrant proteins (Balch et al., 2008; Clausen et al., 2019; Hartl et al., 2011). During translation, Hsp70 and other molecular chaperones promote the folding of nascent proteins, but also assist the refolding of proteins that become damaged after synthesis (Hartl and Hayer-Hartl, 2009). Conversely, degradative PQC typically relies on the ubiquitin-proteasome system (UPS) to eradicate non-native proteins from the intracellular space (Clausen et al., 2019; Enam et al., 2018). Here, often assisted by molecular chaperones, non-native proteins are conjugated to ubiquitin-chains by E3 ubiquitin-protein ligases and targeted to the 26S proteasome for degradation.

Virtually all proteins may become structurally destabilized. The PQC system must therefore have a wide-ranging substrate specificity to ensure that any destabilized protein can be recognized, while still safeguarding that native proteins are not targeted. Despite several efforts (Geffen et al., 2016; Maurer et al., 2016; Rosenbaum et al., 2011; van der Lee et al., 2014), the discerning features in a destabilized protein, the so-called PQC degrons (Ravid and Hochstrasser, 2008) that are recognized by E3s and thus trigger degradation, still remain rather enigmatic. These degrons are, however, expected to include hydrophobic areas that are buried in the native protein but become exposed upon misfolding (Ravid and Hochstrasser, 2008).

To further unravel the nature of the PQC system, we investigated the degradation of human dihydrofolate reductase (DHFR), a known PQC target in *E. coli* (Thompson et al., 2020) and where certain germline missense variants have been linked to recessive megaloblastic anemia (Banka et al., 2011; Cario et al., 2011). We show that the disease-linked missense variants structurally destabilize the DHFR protein, which in turn is degraded by the UPS. We identified two DHFR regions, DHFR-Deg13-36 and DHFR-Deg61-84 (abbreviated Deg1 and Deg2, respectively), which display degron properties, and find that Deg2 degradation requires Hsp70. Deg2 is positioned in a buried region of DHFR, but H/D exchange studies revealed that the backbone amide hydrogens of Deg2 are more prone to exchange with solvent hydrogens in the disease-linked DHFR than in the wild type, suggesting that it operates to trigger degradation upon partial unfolding.

## Results

### Disease-linked DHFR missense variants are structurally destabilized and rapidly degraded

Clinical genetics studies have shown that two *DHFR* missense variants (L80F and D153V) are linked to megaloblastic anemia (OMIM: 613839) (Banka et al., 2011; Cario et al., 2011). In both cases the authors found that the protein variants displayed reduced steady-state levels (Banka et al., 2011; Cario et al., 2011), suggesting that the proteins are PQC targets. To test this in a genetically tractable model organism, we expressed wild-type human DHFR and the two disease-linked variants in yeast cells and compared the levels by Western blotting. As expected, both L80F and D153V were present at reduced levels compared to wild-type DHFR (Fig. 1A). This reduced level was caused by proteasomal degradation since the protein levels were, in both cases, restored when the proteasome was blocked by bortezomib (BZ) (Fig. 1B). This suggests that the disease-linked DHFR proteins are structurally destabilized PQC targets. However, since addition of the DHFR inhibitor methotrexate (MTX) also led to a stabilization of the DHFR variants (Fig. 1C), the structural destabilization is not so severe that the DHFR variants are unable to bind MTX. Accordingly, both variants were soluble cytosolic proteins (Fig S1A), adept at binding a MTX-resin (Fig. S1B), and conferred MTX-resistance to the yeast cells when overexpressed, albeit not to same extent as wild-type DHFR (Fig. S1C). In agreement with the cellular studies, we found that both purified DHFR variants in their folate bound states displayed at least 6 °C reduction in the unfolding midpoint from the heat denaturation experiment compared with folate bound wild-type DHFR (Fig. 1D). The temperature unfolding midpoints are 43.5 °C and 45.1 °C for L80F and D153V, respectively, compared to 51.1 °C for wild-type DHFR. We also note that the temperature unfolding curves in the presence of folate are rather shallow and that around 10% of the protein is already unfolded at 37 °C for the variants. When DHFR is bound to MTX, there is a clear increase in the unfolding temperature of both wild type and the variants. The two variants are only destabilized by approximately 3 °C in the presence of MTX and the unfolding curves become steeper than those recorded in the presence of folate. Thus, the disease-causing mutations significantly destabilize the folded structure of DHFR, but the destabilization may be alleviated by binding of ligands. Indeed, we were unable to purify the DHFR variants in the ligand-free apo-state.

**Fig. 1.**
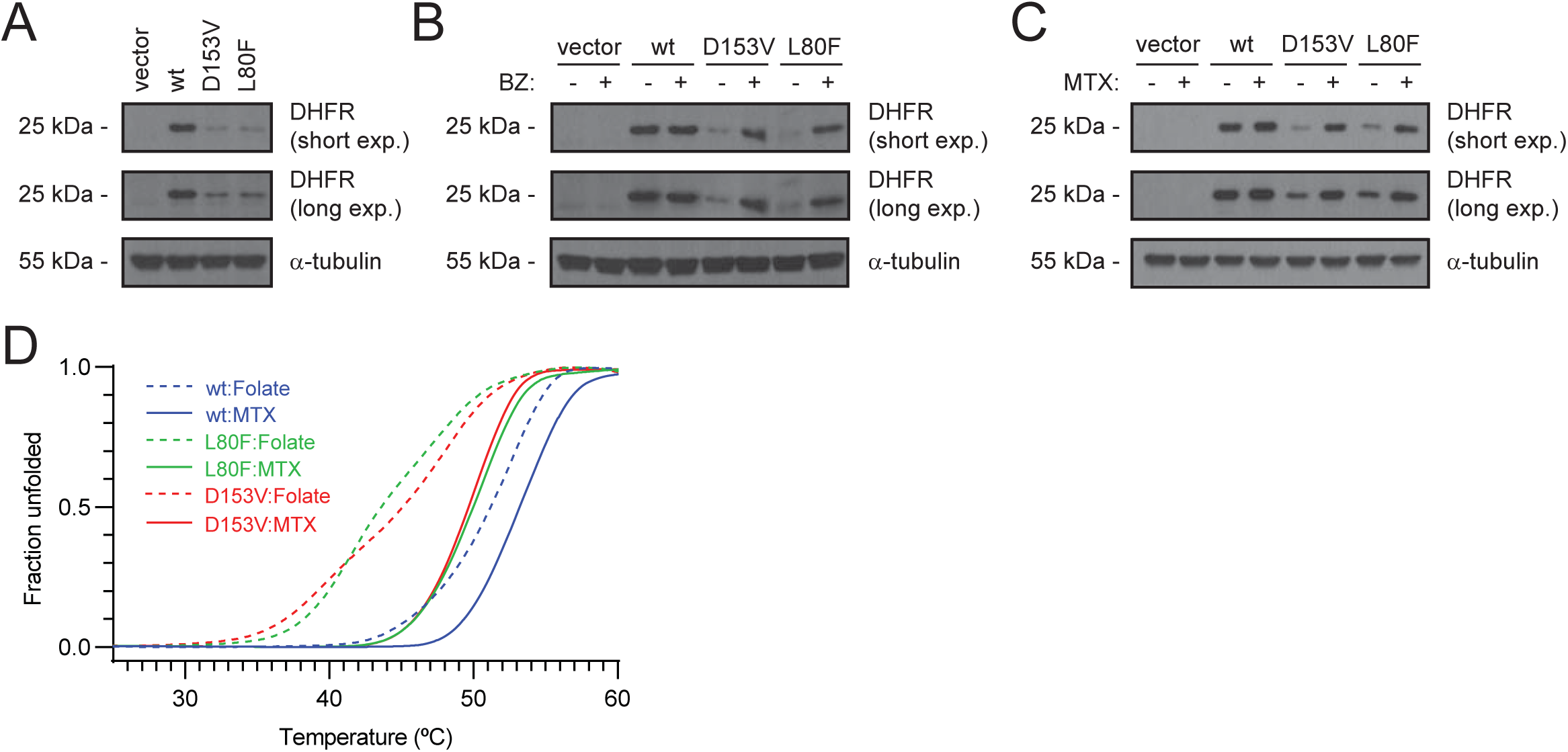
The disease-linked DHFR variants are functional but unstable proteasome targets. (A) The steady-state levels of the DHFR variants in wild-type cells were compared by SDS-PAGE and Western blotting using antibodies to DHFR. α-tubulin served as a loading control. (B) Wild-type cells transformed to express the indicated DHFR variants were grown with (+) or without (-) the proteasome inhibitor bortezomib (BZ), and cell lysates were compared by SDS-PAGE and Western blotting using antibodies to DHFR. α-tubulin served as a loading control. (C) Wild-type cells transformed to express the indicated DHFR variants were grown with (+) or without (-) the DHFR inhibitor methotrexate (MTX), and cell lysates were compared by SDS-PAGE and Western blotting using antibodies to DHFR. α-tubulin served as a loading control. (D) Heat denaturation of DHFR variants with either folate or methotrexate bound. The unfolding was followed by intrinsic Trp fluorescence at 350 nm and the signal normalized to give the fraction of unfolded protein.

### Systematic screen for PQC degrons in DHFR

Since the above data show that the disease-linked DHFR protein variants are proteasome targets, we proceeded to map potential degrons in DHFR using a yeast selection system (Geffen et al., 2016) where potential degrons are fused in-frame downstream of Ura3-HA-GFP fusion protein (Fig. 2A). Thus, in case a fragment containing a degron is fused to the reporter, the processive nature of proteasomal degradation ensures that the entire fusion protein is degraded and cells carrying a deletion in *ura3* should therefore not be able to grow unless the medium is supplemented with uracil (Fig. 2A).

**Fig. 2.**
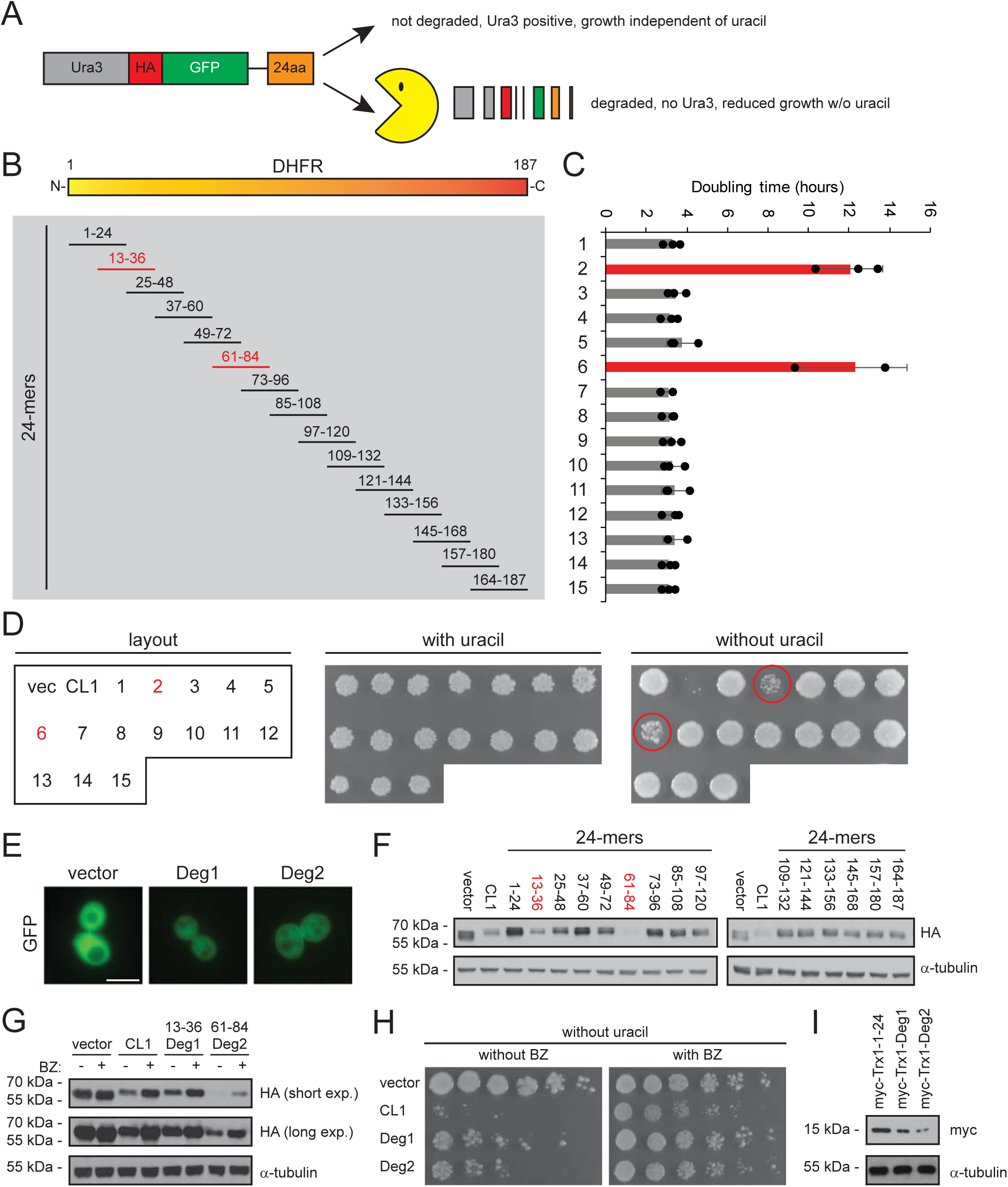
Fragments of the human DHFR protein display degron activity. (A) In the yeast selection system used for identifying degrons, the different 24-residue DHFR fragments are fused after a linker to the C-terminus of a Ura3-HA-GFP fusion protein. In case a fragment acts as a degron, the entire fusion protein will be degraded and the cells will be unable to grow without addition of uracil to the medium. (B) The 187-residue human DHFR protein (yellow/orange bar) was divided into the indicated 15 different 24-residue partially overlapping fragments (24-mers) and fused to the C-terminus of the Ura3-reporter. (C) Growth was monitored in medium lacking uracil and the doubling times were determined (right panel). Two fragments (red) led to strongly reduced growth (increased doubling times). The vector and CL1 fusion protein were included as controls. The error bars indicate the standard deviation (n=3). (D) The growth of yeast cells containing the indicated fragments (left panel and panel B) was monitored by spotting an equal amount of culture onto solid media with (central panel) or without (right panel) uracil. Note the fragments displaying increased doubling times in panel C also grow poorly on solid medium without uracil (circled). The vector and CL1 fusion protein were included as controls. (E) Cells carrying the indicated expression constructs were analyzed by fluorescence microscopy. The Deg1 and Deg2 fusion proteins are cytosolic and present in reduced amounts compared to the vector control. Bar = 5 µm (F) The steady-state levels of cells carrying the indicated expression constructs were analyzed by SDS-PAGE and Western blotting using antibodies to the HA-tag on the reporter protein. α-tubulin served as a loading control. (G) The vector, CL1, Deg1, and Deg2 strains were grown with (+) or without (-) the proteasome inhibitor bortezomib (BZ), and whole cell lysates were compared by SDS-PAGE and Western blotting using antibodies to the HA-tag on the reporter fusion. α-tubulin served as a loading control. (H) Growth was compared on solid medium without uracil and with or without bortezomib (BZ), by spotting serial dilutions of the indicated cultures. (I) The steady-state amounts of myc-tagged Trx1 fused in frame to the indicated fragments of DHFR were compared by SDS-PAGE and Western blotting of cell extracts using antibodies to the myc tag. α-tubulin served as a loading control.

The 187 residue full-length DHFR protein was divided into 15 different partially overlapping 24-residue fragments and produced in frame with the Ura3-reporter (Fig. 2AB). The artificial 16-residue CL1 peptide that is a known degron (Gilon et al., 1998; Gilon et al., 2000; Metzger et al., 2008; Ravid et al., 2006) was included as a control. The length of 24 residues was selected since we reasoned that fragments of this size are likely sufficiently long to contain degradation signals, but too short to harbor significant structure. The growth of most strains was unaffected by the C-terminal fusions (Fig. 2BC), suggesting that these fragments do not contain degrons. However, fragments composed of DHFR residues 13-36 and residues 61-84 led to significantly reduced growth in both liquid growth assays (Fig. 2C) and on solid medium (Fig. 2D). From here on, we will refer to the 13-36 fragment as Deg1 and the 61-84 fragment as Deg2.

Fluorescence microscopy revealed that the Deg1 and Deg2 fusions were cytosolic (Fig. 2E). However, in agreement with the degrons leading to reduced cellular amounts of the reporter protein, fusion of the degrons to the reporter led to clearly reduced GFP intensities (Fig. 2E). This was also evident in Western blots of whole cell lysates (Fig. 2F). The reduced amount of the degron fusion proteins was caused by proteasomal degradation, since adding sublethal amounts of BZ led to increased amounts of the reporter proteins (Fig. 2G), and also restored cell growth in the absence of uracil (Fig. 2H).

Finally, to test if the degron properties of Deg1 and Deg2 were transferrable, we generated fusions between the cytosolic thioredoxin, Trx1, and Deg1, Deg2 and, as a control, the 1-24 fragment of DHFR that did not appear destabilizing in context of the Ura3 fusion (Fig. 2CDF). We observed that the Deg1 and Deg2 fusions with Trx1 led to reduced protein levels (Fig. 2I).

Considering that relevant PQC degrons should be exclusively found in buried regions, we mapped the position of Deg1 and Deg2 on the DHFR structure (PDB: 1U72) (Cody et al., 2005). Both degrons contained exposed and buried parts (Fig. 3AB). However, when, as a measure of exposure, we mapped the number of C-beta neighbors (C-alpha for glycine) within 8 Å of each residue, the central region of Deg2 clearly contains a buried stretch of residues (Fig. 3C & supplemental file 1). To probe these sequences further, we generated a number of N- and C-terminal truncations of Deg2 (Fig. 3D) and tested their degron activity as before (Fig. 3EF). Growth assays revealed a minimum 8-amino acid residue sequence (RINLVLSR) that was required for degron activity and led to reduced levels of the reporter protein (Fig. 3EF). This central Deg2 region bears some resemblance to the preferred Hsp70-binding sequence consisting of hydrophobic residues flanked by positive charges (Rudiger et al., 1997). We therefore proceeded to test if the degron activity required Hsp70. In the case of Deg1, we only observed very slight effects (Fig. S2AB), but for Deg2, the degron activity was clearly dependent on the Hsp70 paralogues Ssa1 and Ssa2 (Fig. S2AB). This suggests that Deg2 indeed represents a PQC degron that upon exposure targets proteins for chaperone-assisted proteasomal degradation. In agreement with this, the stability of the full-length DHFR variants was also increased in mutants lacking Ssa1 or Ssa2 (Fig. S2C).

**Fig. 3.**
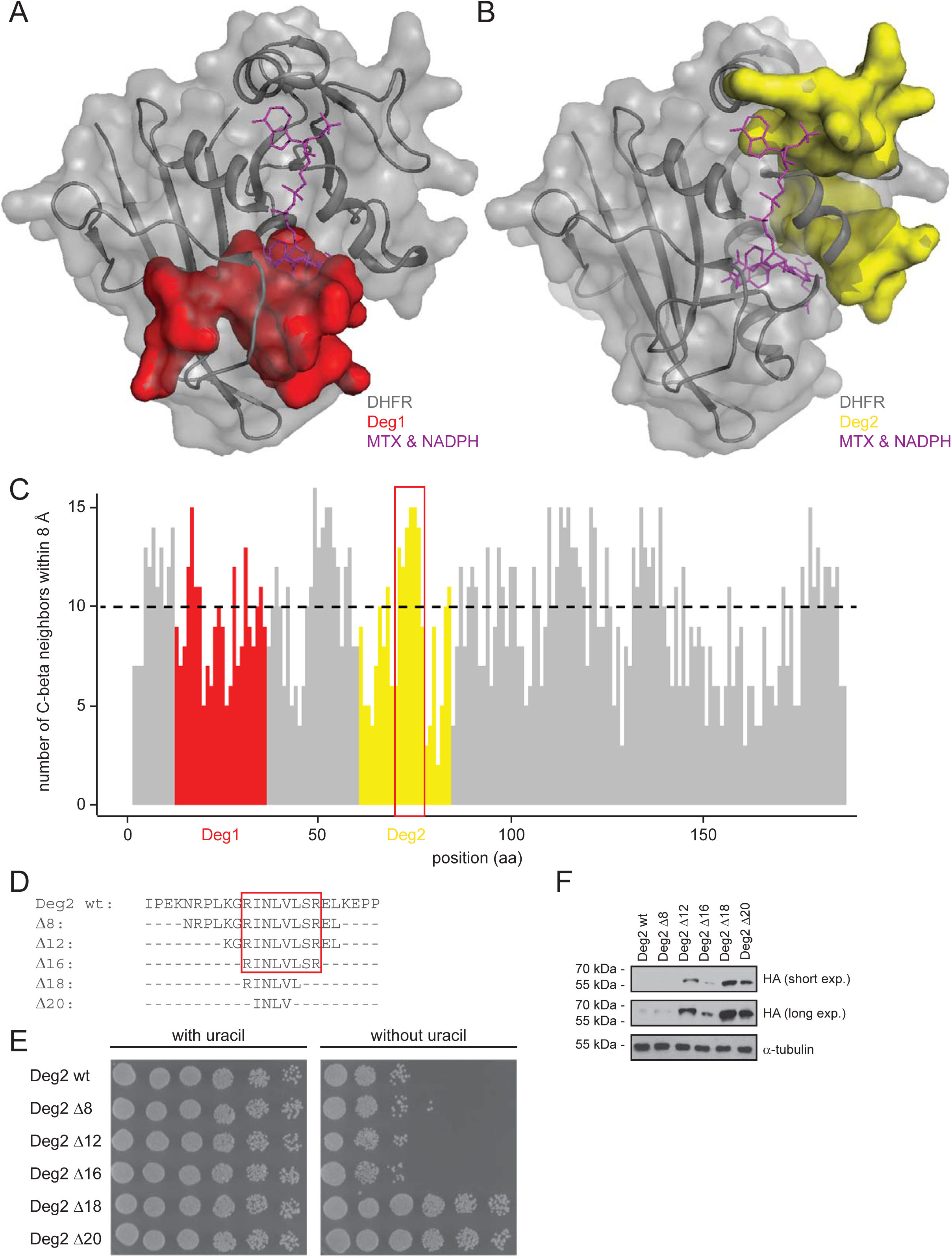
Positioning of Deg1 and Deg2 in DHFR. (A) The position of Deg1 in the DHFR structure (PDB: 1U72). Deg1 is marked in red. The inhibitor and substrate methotrexate (MTX) and NADPH, respectively, are shown in purple. (B) The position of Deg2 in the DHFR structure. Deg2 is marked in yellow, while methotrexate (MTX) and NADPH are purple. (C) The number of C-beta neighbors for each residue in DHFR is shown. Deg1 is shown in red. Deg2 is shown in yellow. The dashed line marks a threshold of 10 neighbors. Note that a high value indicates that the residue is buried. The red box marks the central buried part of Deg2. (D) The amino acid sequence of the tested truncations of the Deg2 fragment. The minimal degron region is boxed. (E) The growth of cells carrying the indicated Deg2 truncations was compared by spotting serial dilutions of the cultures on solid medium with or without uracil. (F) The steady-state amounts of the indicated Deg2 variants were compared by SDS-PAGE and Western blotting of whole cell extracts using antibodies to the HA tag. α-tubulin served as a loading control.

### H/D-exchange NMR

To test if the identified degron regions in DHFR were relevant in context of the full-length DHFR protein and disease-linked protein variants, we next decided to measure the local structural stabilities by hydrogen-deuterium exchange of the backbone amides by NMR spectroscopy. HSQC spectra show that the structures of the DHFR variants overall are the same as for the wild-type, but that there are some local differences around the mutation sites. At the first time point in the hydrogen exchange time series, recorded approximately five minutes after the lyophilized protein was dissolved in D_2_O, we observed peaks from 129, 122, and 125 of the 174 non-proline backbone amides in wild-type, L80F and D153V, respectively (Fig. 4A). Within the 5352 min (3.7 days) for which the hydrogen exchange was followed, we could quantify the exchange kinetics for most of the observed amide residues. For some residues the exchange was either too slow or too fast to be quantified (Fig. S3). We thus quantified the change in the exchange rates for 97 of the amides in the two disease-linked variants relative to the wild-type (Fig. 4B). Most amides exchange faster in the two variants, although in both variants some also exchange slower. These slower exchanging residues are probably a result of subtle structural changes induced by the mutations that stabilize isolated hydrogen bonds. In L80F, we observe faster exchange kinetics mostly between residue 51 and 126, which corresponds to the residues in the vicinity of position 80. The L80F mutation thus results in local destabilization of the hydrogen bonds, including those involving the amides in Deg2 (Fig. 4C). However, L80F appears to leave the local stability of the Deg1 region unaffected. D153V on the other hand has a more global effect on the exchange rates and affects residues throughout the structure (Fig. 4C), and the hydrogen bonds in both Deg1 and Deg2 are weakened. The increase in the exchange rates for both variants are between 2^1^ and 2^4^, which corresponds to an amide in the variants on average spends 2 to 16 times more time in the open state than the same amide in wild-type does. This estimate assumes that the exchange process follows the EX2 limiting case of the Linderstrøm-Lang model for hydrogen exchange which dominates at neutral and low pH (Hvidt and Nielsen, 1966).

**Fig. 4.**
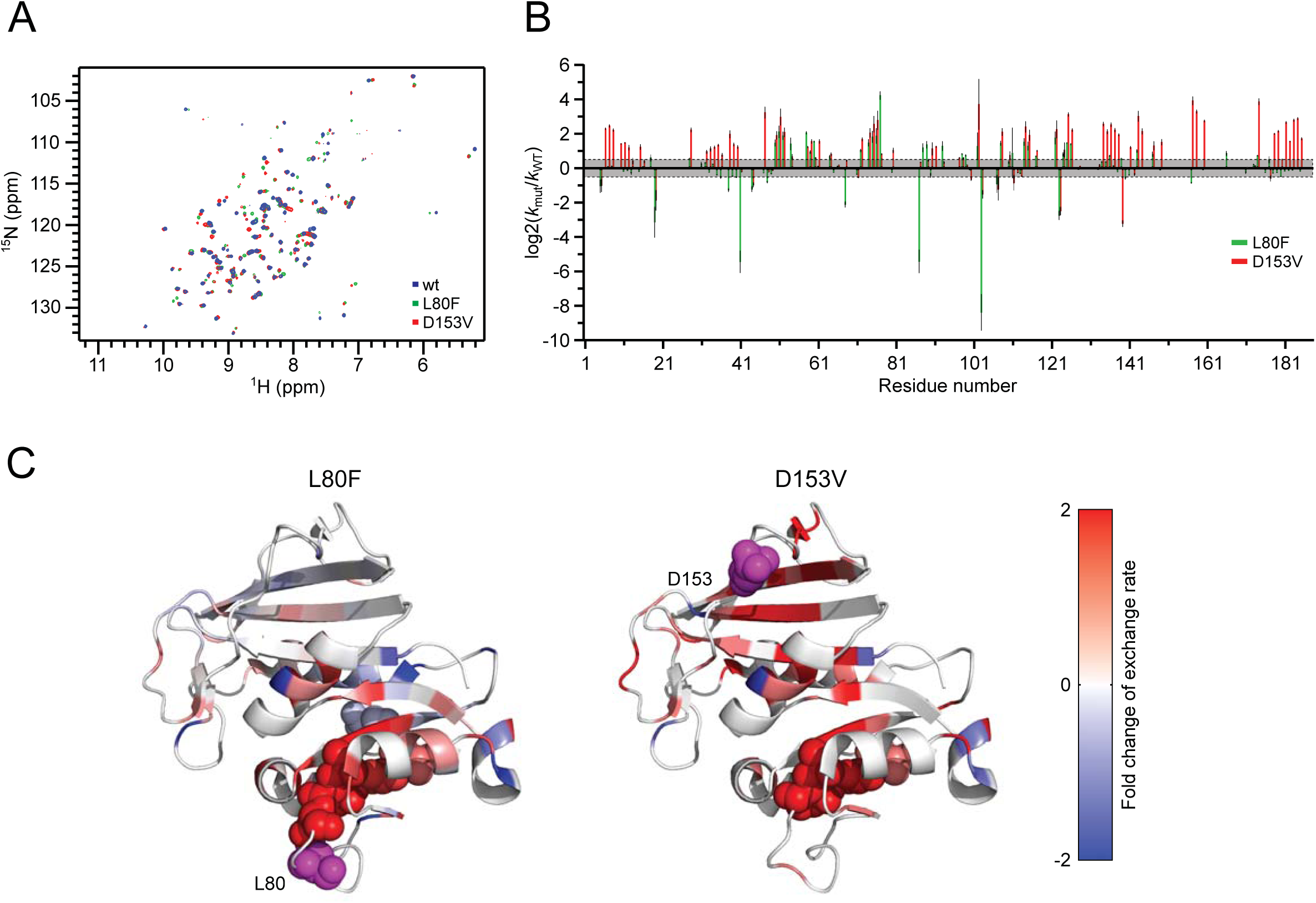
Local stability change in DHFR variants from hydrogen deuterium exchange. (A) First ^15^N-HSQC spectrum from the HDX series of wild-type (blue), L80F (green) and D153V (red). (B) Residue specific change in the measured hydrogen exchange rate, *k*_mut_, for the two mutants relative to the exchange rate for the wild-type, *k*_wt_. Log base 2 of the ratio is plotted and shows how many times the hydrogen exchange rate of the mutants have doubled relative to the wild-type. (C) The values in (B) plotted on the crystal structure of DHFR with a color scale from blue (−2) to red (2). The mutation sites are shown as magenta spheres. The residues of Deg2 for which we have HDX information are also shown as spheres color-coded as the rest of the structure.

In conclusion, our hydrogen-deuterium exchange data show that the disease-linked DHFR variants more frequently visit a locally unfolded state where the hydrogen-bonds in the eight-residue segment comprising Deg2, are broken.

## Discussion

Currently, it is well established that certain missense mutations may cause protein destabilization and degradation (Stein et al., 2019), and that this is the underlying molecular mechanism for a number of hereditary diseases, including cystic fibrosis, phenylketonuria and various cancer predisposition disorders (Abildgaard et al., 2019; Ahner et al., 2007; Arlow et al., 2013; Canaff et al., 2012; Clausen et al., 2020; Gersing et al., 2021; Pey et al., 2007; Scheller et al., 2019). Here, we add hereditary megaloblastic anemia, linked to germline *DHFR* variants, to this list. Our results reveal that both DHFR protein variants are PQC targets that are structurally destabilized and degraded. However, upon binding methotrexate, the proteins are significantly stabilized and no-longer PQC targets. This suggests that the DHFR variants are not strongly misfolded, and still adept at binding MTX. These observations are in line with previous experiments using a destabilized, aggregation prone variant of *E. coli* DHFR, which showed that this variant could be stabilized inside *E. coli* by addition of the DHFR ligand trimethoprim (Hingorani et al., 2017). In the case of the L80F and D153V variants of human DHFR, we speculate that they are likely to be partially functional, given that *DHFR* is an essential gene, and disease-linked *DHFR* variants are thus bound to be hypomorph rather than null alleles.

Although several regions of DHFR become more dynamic in the disease variants, one region in particular overlaps with a mapped degron region, namely an 8-residue (RINLVLSR) buried fragment from position 71-78. Importantly, this region, similar to the full-length DHFR variants, requires Hsp70 for degradation, while the other identified degron appears to function independently of Hsp70. This suggests that upon the mutation-induced structural destabilization, there is increased transient exposure of Deg2, which in turn leads to chaperone-dependent proteasomal degradation of the DHFR variants. This is in agreement with a number of other studies that have linked molecular chaperones with the UPS (Gowda et al., 2013; Kampmeyer et al., 2017; Kandasamy and Andreasson, 2018; Kriegenburg et al., 2014; Mathiassen et al., 2015; Samant et al., 2018), including as E3 co-factors involved in substrate selection (Murata et al., 2001; Qian et al., 2006; Xu et al., 2002), or as carriers of misfolded proteins to E3 ligases (Comyn et al., 2016; Guerriero et al., 2013; Shiber and Ravid, 2014). For instance, the human E3 ligase, CHIP, associates directly with Hsp90 and Hsp70 to catalyze the ubiquitylation of chaperone-bound clients (Connell et al., 2001; Demand et al., 2001). We thus propose that Deg2 functions as a classical PQC degron, in that it is buried in wild-type DHFR, but becomes exposed and elicits chaperone-dependent degradation in the destabilized disease-linked DHFR variants. Deg1, on the other hand, seems irrelevant for the tested DHFR variants. Interestingly, in comparison with Deg2, Deg1 appears more exposed, suggesting that in the native protein, Deg1 is maintained in a conformation that does not trigger degradation. As a 24-mer peptide, however, and presumably in the fully denatured state of DHFR, it would become recognized by an E3. The two DHFR variants that we tested here are currently the only known disease-linked alleles of DHFR. However, it is possible that Deg1 is relevant for other, presently unknown DHFR variants, that may become structurally perturbed in a way leading to Deg1 recognition.

In conclusion, the results presented here provide detailed mechanistic insight into the degradation of disease-relevant PQC targets and show an example of how a hydrophobic and buried region in a native protein, may become exposed and target the protein for proteasomal degradation.

## Materials and methods

### Plasmids

Human *DHFR* cDNA fragments were inserted into the pTR1412 vector (Geffen et al., 2016) in frame after the *URA3-HA-GFP* fusion (Genscript). Full-length *DHFR* was expressed in yeast from pTR1412 lacking the *URA3-HA-GFP* fusion (Genscript). Point mutations were generated by Genscript. For expression of the Trx1 fusion proteins, the fragments were inserted in frame after myc-tagged *TRX1* in the pESC vector (Genscript). For production of DHFR and DHFR variants in *E. coli*, codon optimized human DHFR was produced as a 6His-SUMO1 fusion from pET28a (Genscript).

### Yeast strains and techniques

All *Saccharomyces cerevisiae* yeast strains were obtained from the Euroscarf collection. For culturing, synthetic complete (SC) medium (2% glucose, 6.7 g/L yeast nitrogen base without amino acids and with ammonium sulfate (Sigma)) supplemented with 1.92 g/L Drop-out Mix Synthetic (US Biological) as required for selection. Yeast cells were transformed using lithium acetate (Gietz and Woods, 2002). For solid media growth assays the strains were cultured at 30°C to exponential phase. The cultures were diluted to an OD_600nm_ of exactly 0.40 and dilution series (5-fold) were prepared. Then 5 µL of each dilution was applied in drops on agar plates. Colonies formed after 2-3 days incubation at 30°C. In growth assays, Bortezomib (LC laboratories) was used at a final concentration of 0.5 mM. For liquid media growth assays, pre-cultures were diluted to an OD_600nm_ of 0.01 and added to a 96-well plate with a cover (Nunc) in triplicates. Wells only containing media, were included as a reference. Turbidity of the cultures was monitored as OD_600nm_ in an Infinite 200 PRO (Tecan) microplate reader for 24 hours at 30°C. The growth curves were used for estimating the doubling times. For SDS-PAGE and Western blotting of the degron-fusions cell lysates were prepared from exponential phase cultures treated with 0.1 mM CuSO_4_ overnight to induce expression. The full-length DHFR variants were produced without CuSO_4_. Bortezomib (LC laboratories) was used at a final concentration of 1 mM and methotrexate at a final concentration of 250 µM. The cells were lysed using glass beads and trichloroacetic acid (TCA) as described (Cox et al., 1997). For microscopy, live cells in exponential phase were imaged using a Zeiss Axio Imager Z1 microscope equipped with a Hamamatsu ORCA-ER digital camera.

### Solubility and co-precipitation experiments

Protein solubility was analyzed from yeast cells in exponential phase. Cell lysates were prepared using glass beads in buffer A (25 mM Tris/HCl pH 7.4, 50 mM NaCl, 10% glycerol, 2 mM MgCl_2_, 1 mM PMSF and Complete Mini Protease Inhibitors (Roche)). The whole cell lysates were centrifuged (13000 g, 30 min.) to separate the soluble and insoluble proteins. The pellet was resuspended in a volume of buffer A matching the volume of the supernatant, then SDS sample buffer was added and the samples were analyzed by SDS-PAGE and Western blotting.

Co-precipitation experiments were performed from exponential phase cultures treated with 0.1 mM CuSO_4_ overnight to induce expression. Cells were lysed using glass beads in 1 cell volume of buffer B (20 mM Tris/HCl pH 7.4, 1 mM EDTA, 1 mM DTT, 10% glycerol, 1 mM PMSF and Complete protease inhibitors (Roche)). The extracts were cleared by centrifugation (13000 g, 30 minutes), and the soluble fraction tumbled end-over-end for 2 hours with 15 µL (bed volume) of methotrexate-agarose (Sigma). Then the beads were washed once in 1 mL of buffer B with 0.5 M KCl, three times with buffer B containing 0.5 M KCl and 1% Triton X-100, and finally once in buffer B with 0.5 M KCl. Protein was eluted in buffer B with 0.5 M KCl and 1 mM methotrexate (Sigma). Then the protein was precipitated with 40% TCA for 60 minutes on ice followed by centrifugation (10000 g, 30 min.). The pellets were extensively washed with ice-cold acetone and air-dried. Finally, the pellets were resuspended in SDS sample buffer and analyzed by SDS-PAGE and Western blotting.

### Electrophoresis and blotting

For SDS-PAGE and Western blotting, protein samples were resolved on 12.5% acrylamide gels and transferred to 0.2 µm nitrocellulose membranes by electro-blotting. Blocking was performed in PBS (10 mM Na_2_HPO_4_, 1.8 mM KH_2_PO_4_, 137 mM NaCl, 3 mM KCl, pH 7.4) with 5% milk powder. The antibodies used were: anti-DHFR (ProteinTech, 15194-1-AP), anti-HA (Roche, clone 3F10), anti-α-tubulin (Abcam, clone TAT1), anti-Pma1 (Abcam, Ab4645), and anti-myc (Chromotek, clone 9E1). The secondary antibodies were from DAKO.

### In silico methods

Protein structures were visualized using PyMOL version 2.3 and the DHFR structure (PDB: 1U72) (Cody et al., 2005). C-beta neighbors were calculated based on the coordinates of PDB 1U72. For each residue, we calculated the number of C-beta atoms within 8 Å of its C-beta (C-alpha for glycine) and set this as the number of C-beta neighbors. Residues with > 10 C-beta neighbors are considered buried, while those with fewer C-beta neighbors are considered exposed.

### Purification of DHFR variants, expressed in E. coli

The DHFR variants were produced in *E. coli* BL21(DE3) pLysS from a pET28a(+)-based plasmid fused N-terminally to a 6His-SUMO tag. Overnight cultures were diluted 200-fold in 1 L M9 minimal media supplemented with 2 mM folic acid, kanamycin and chloramphenicol, and incubated at 37 °C. For single and double-labelled protein production, the NH_4_Cl and glucose content of the M9 minimal media were replaced with ^15^NH_4_Cl (CortecNet) and/or ^13^C-glucose (CortecNet), respectively. When OD_600nm_ reached 0.5, the cultures were moved to 15 °C for 30 minutes before addition of IPTG to 0.5 mM. After 16-18 hours the cells were harvested by centrifugation. The cell pellet was resuspended in 40 mL lysis buffer (50 mM NaH_2_PO_4_ pH 7.4, 300 mM NaCl, 1 mM PMSF and Complete protease inhibitors without EDTA (Roche)) and sonicated on ice. The lysate was cleared by centrifugation (13000 g, 30 min) and the supernatant passed over a TALON resin (Clontech) and purified according to the manufacturer’s instructions. Fractions containing the DHFR protein, were pooled and dialyzed against cleavage buffer (25 mM Na-phosphate pH 7.4, 150 mM NaCl, 0.5 mM folate) overnight at 4 °C. The following day, DTT was added to a final concentration of 0.5 mM and His-SUMO protease (Sigma) was added to a concentration 100-fold lower than the total protein concentration. After 2 hours at 4 °C the reaction mixture was passed over a second TALON resin column, and the cleaved DHFR was collected as flow-through. Pooled fractions were dialyzed overnight against NMR buffer (50 mM K-phosphate pH 6.5, 100 mM KCl, 1 mM DTT, 25 µM folic acid). By SDS-PAGE analysis the purified proteins were estimated to be >95% pure. The protein concentration was determined by a bicinchoninic acid (BCA) assay (Pierce) using bovine serum albumin (BSA) as a standard. The yield was roughly 20-30 mg of DHFR from 1 L of M9 culture.

### Heat denaturation

Samples of 25 μM of each DHFR variant were prepared by dilution in 50 mM K-phosphate, 100 mM KCl, 1 mM DTT, 25 µM folate, pH 6.5. To replace bound folate, 50 μM MTX was added to the folate samples. The unfolding of DHFR was followed by intrinsic Trp fluorescence using a Nanotemper Prometheus NT.48 fluorimeter. Samples of 15 μL were placed in high sensitivity quarts capillaries (Nanotemper) and the temperature was ramped with 1 °C/min from 20 °C to 95 °C. The fluorescence signal at 350 nm was normalized to represent the fraction of unfolded protein.

### Chemical shift assignments

^13^C-^15^N double labelled samples of all variants were prepared with a protein concentration of 300 µM in 50 mM K-phosphate, 100 mM KCl, 1 mM DTT, 25 µM folate, pH 6.5. ^15^N-HSQC, HNCO, HNCA, HNCOCA, HNCOCACB and HNCACB were recorded on a 800 MHz Bruker Avance III spectrometer at 25 °C. The triple resonance spectra were recorded with 25% non-uniform sampling and reconstructed by compressed sensing using the iterative soft threshold algorithm with virtual echo implemented in qMDD (Kazimierczuk and Orekhov, 2011) and processed using nmrPipe (Delaglio et al., 1995). Backbone H^N 15^N^H, 13^C’, ^13^Cα and ^13^Cβ chemical shifts were assigned using ccpnmr analysis (Skinner et al., 2016) and the previously published assignment of wild-type DHFR (Bhabha et al., 2013).

### Hydrogen-deuterium exchange

To measure the hydrogen exchange in D_2_O, 600 µL of 400 µM ^15^N-labelled protein in 50 mM K-phosphate, 50 mM KCl, 1 mM DTT, 25 µM folate, pH 6.5 were lyophilized, and the resulting protein powder resuspended in 600 µL cold 99.9% D_2_O. The HDX time series of all three variants were recorded on a 750 MHz Bruker Advance III HD spectrometer at 25 °C using a standard HSQC pulse program. The acquisition time for each experiment was 19.53 minutes and 275 HSQC were recorded for each variant corresponding to 91.5 hours.

The spectra were processed, and peaks intensities measured with nmrPipe (Delaglio et al., 1995). For each peak, the data was fitted to a single exponential decay with an offset, I(t) = (I_o_-I_∞_)*exp(-*k*\*t) + I_∞_. For amides where the exchange reached a final plateau in all three variants, I_∞_ was fitted as a global parameter for all three variants. For residues where a plateau was not reached, the data were fitted with I_∞_ fixed at the average value from the residues where this parameter could be estimated.

## Acknowledgements

The authors thank Anne-Marie Lauridsen for technical assistance, and Dr. Michael Askvad Sørensen for assistance with DHFR purification.

## Conflict of interest

No conflicting interests declared.

## Funding

The present was funded by the Novo Nordisk Foundation (https://novonordiskfonden.dk) challenge program PRISM (to K.L.L., A.S. & R.H.P.), NNF18OC0052441 (to R.H.P.) and NNF18OC0032996 (to K.T.), the Lundbeck Foundation (https://www.lundbeckfonden.com) R272-2017-452 and R209-2015-3283 (to A.S.), and Danish Council for Independent Research (Det Frie Forskningsråd) (https://dff.dk) 7014-00039B (to R.H.P.). The funders had no role in study design, data collection and analysis, decision to publish, or preparation of the manuscript.

## Author contributions

C.K., M.M., S.L.-L., A.C., M.R.W., S.V.N., S.L., H.K.M.I. and A.S. performed the experiments, C.K., T.R., A.S., K.L-L., K.T. and R.H.-P. analyzed the data. T.R., K.L.-L. and R.H.-P. conceived the study. C.K. and R.H.-P. wrote the paper.

## Figure legends

**Fig. S1.**
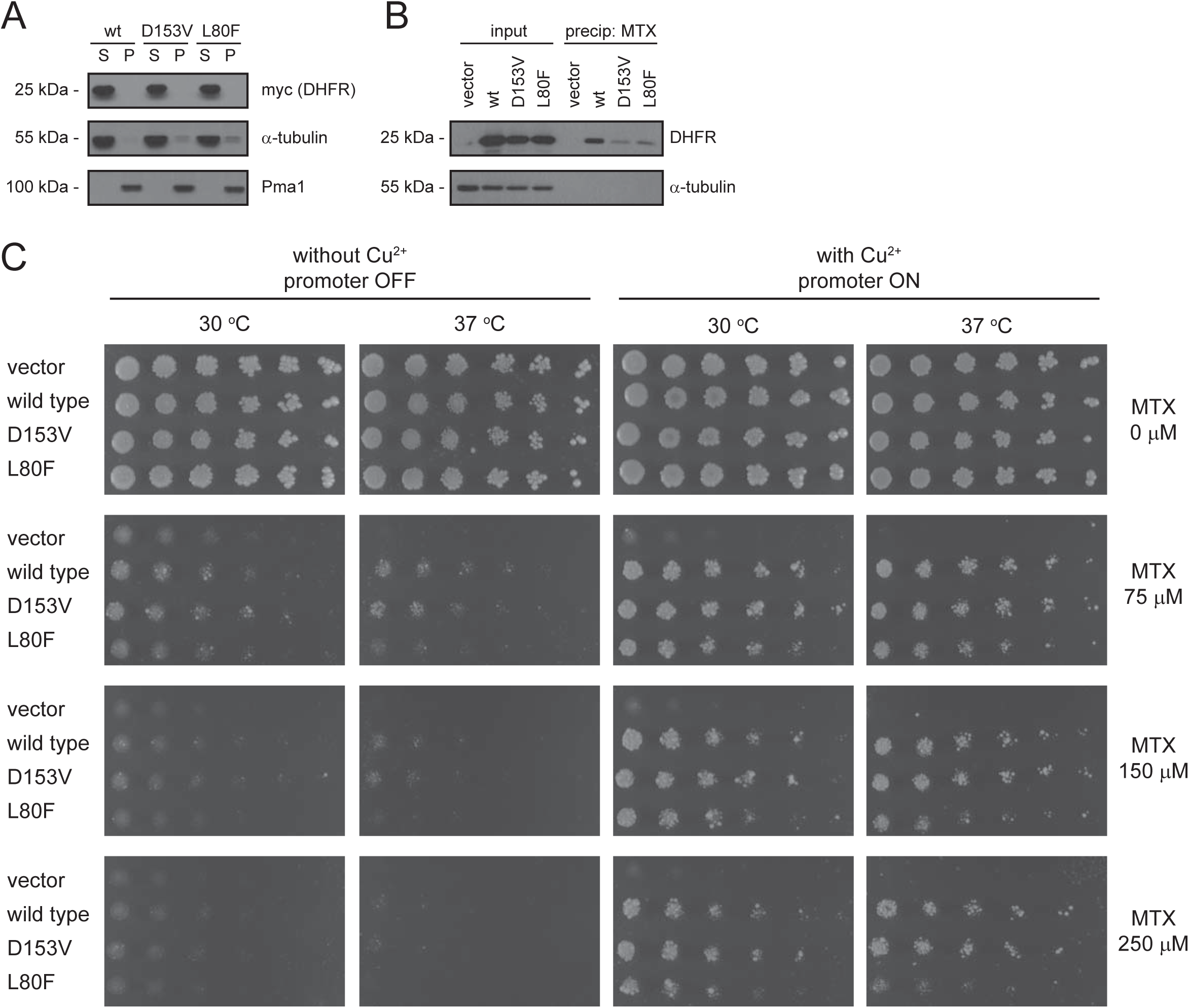
The disease-linked DHFR variants are soluble and bind methotrexate. (A) Cell lysates of wild-type yeast cells expressing the indicated DHFR variants were separated into a soluble supernatant (S) fraction and insoluble pellet (P) fraction by centrifugation. The protein levels in the fractions were compared by SDS-PAGE and Western blotting using antibodies to the myc-tag on DHFR. Blotting for α-tubulin and Pma1 served as controls for a soluble and insoluble protein, respectively. (B) Cleared lysates of wild-type yeast cells transformed to express the indicated DHFR variants (input) were incubated with methotrexate–resin (MTX). After washing, bound proteins were eluted with MTX and analyzed by SDS-PAGE and blotting using antibodies to DHFR. α-tubulin served as a control. (C) The growth of wild-type cells transformed with the indicated DHFR variants was compared at different concentrations of methotrexate (MTX) at 30 °C and 37 °C. Note that DHFR expression was regulated by the Cu-inducible *CUP1* promoter, and accordingly the MTX-resistant phenotype is only evident in the presence of Cu^2+^.

**Fig. S2.**
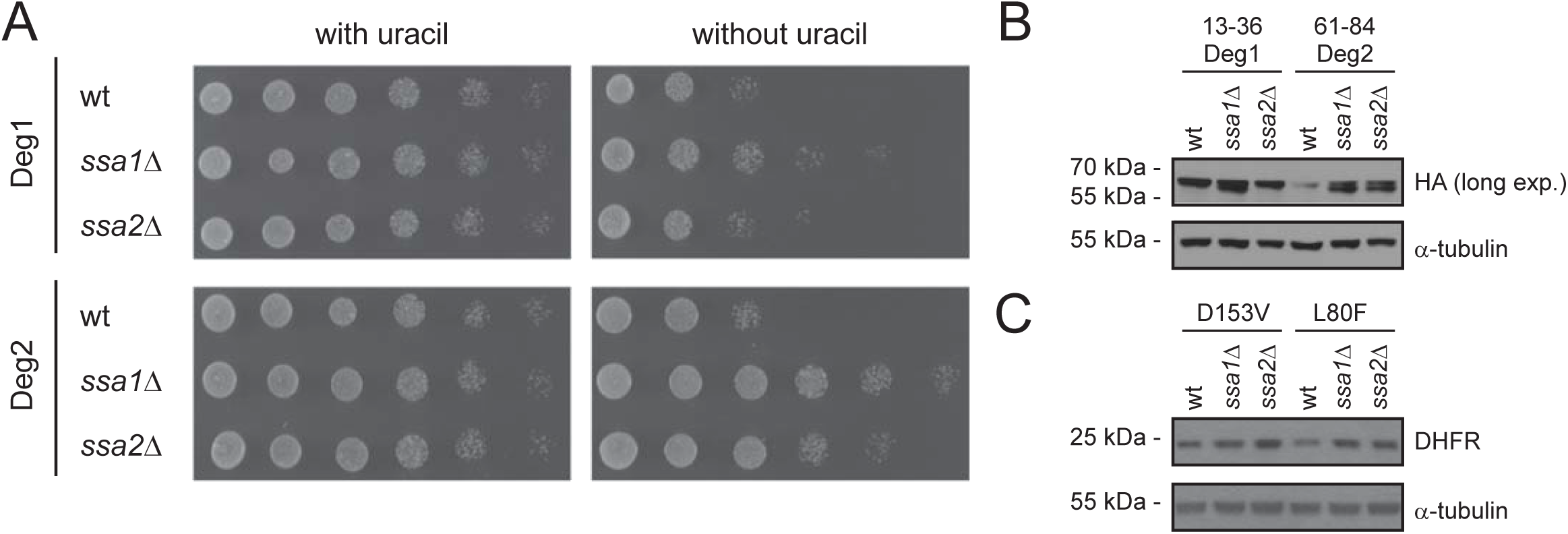
Deg2 and the DHFR variants are targeted by Hsp70 chaperones. (A) The growth of wild type (wt), *ssa1*Δ or *ssa2*Δ cells carrying either the Deg1 or Deg2 constructs as indicated was compared by spotting serial dilutions of the cultures on solid medium with or without uracil. (B) The steady-state amounts of the Deg1 and Deg2 fusion proteins in wild type (wt), *ssa1*Δ and *ssa2*Δ cells were compared by SDS-PAGE and Western blotting of cell extracts using antibodies to the HA tag. α-tubulin served as a loading control. (C) The steady-state amounts of the D153V and L80F DHFR variants in wild type (wt), *ssa1*Δ and *ssa2*Δ cells were compared by SDS-PAGE and Western blotting of whole cell extracts using antibodies to DHFR. α-tubulin served as a loading control.

**Fig. S3.**
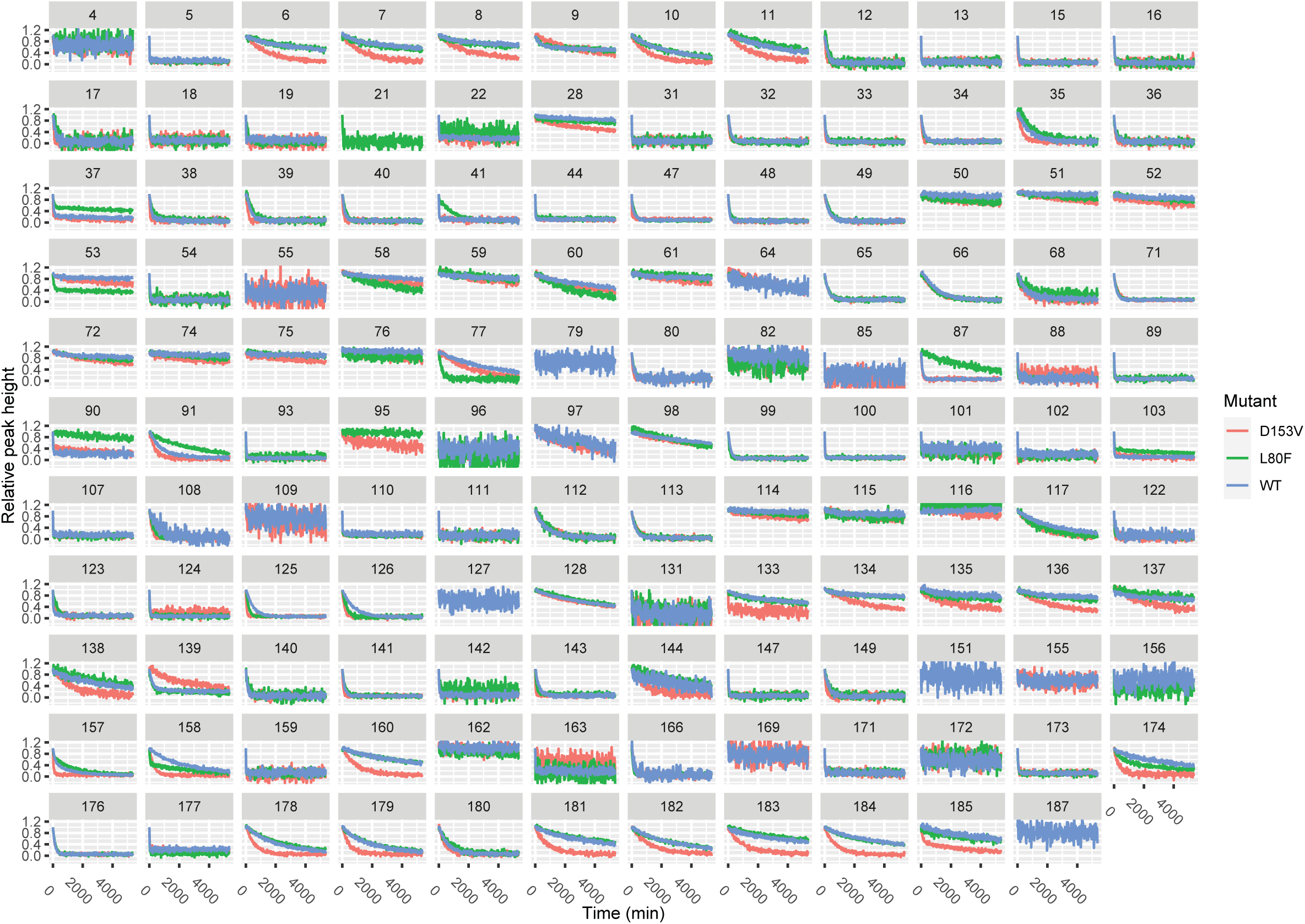
Raw data from the hydrogen deuterium experiment. For each residue observed in first ^15^N-HSQC in the time series, the relative peak intensities are plotted as a function of time.

## Notes

### Competing Interest Statement

The authors have declared no competing interest.

