## SupplementalFile1 for "Disease-linked mutations trigger exposure of a protein quality control degron in the DHFR protein"

Deg1 - red
Deg2 - yellow

| Residue | PDB | EnsEMBL | num.cb.nbrs | cb.nbr.cat | degron | degron.window.1 | degron.window.2 |
|---|---|---|---|---|---|---|---|
| 1 | - | M | NA | NA | no | 0 | 1 |
| 2 | V | V | 7 | exposed | no | 0 | 1 |
| 3 | G | G | 7 | exposed | no | 0 | 1 |
| 4 | S | S | 7 | exposed | no | 0 | 1 |
| 5 | L | L | 13 | buried | no | 0 | 1 |
| 6 | N | N | 12 | buried | no | 0 | 1 |
| 7 | C | C | 14 | buried | no | 0 | 1 |
| 8 | I | I | 11 | buried | no | 0 | 1 |
| 9 | V | V | 13 | buried | no | 0 | 1 |
| 10 | A | A | 10 | exposed | no | 0 | 1 |
| 11 | V | V | 12 | buried | no | 0 | 1 |
| 12 | S | S | 14 | buried | no | 0 | 1 |
| 13 | Q | Q | 9 | exposed | deg1 | 0 | 3 |
| 14 | N | N | 7 | exposed | deg1 | 0 | 3 |
| 15 | M | M | 8 | exposed | deg1 | 0 | 3 |
| 16 | G | G | 12 | buried | deg1 | 0 | 3 |
| 17 | I | I | 15 | buried | deg1 | 0 | 3 |
| 18 | G | G | 11 | buried | deg1 | 0 | 3 |
| 19 | K | K | 11 | buried | deg1 | 0 | 3 |
| 20 | N | N | 5 | exposed | deg1 | 0 | 3 |
| 21 | G | G | 7 | exposed | deg1 | 0 | 3 |
| 22 | D | D | 6 | exposed | deg1 | 0 | 3 |
| 23 | L | L | 9 | exposed | deg1 | 0 | 3 |
| 24 | P | P | 10 | exposed | deg1 | 0 | 3 |
| 25 | W | W | 9 | exposed | deg1 | 2 | 3 |
| 26 | P | P | 5 | exposed | deg1 | 2 | 3 |
| 27 | P | P | 6 | exposed | deg1 | 2 | 3 |
| 28 | L | L | 12 | buried | deg1 | 2 | 3 |
| 29 | R | R | 7 | exposed | deg1 | 2 | 3 |
| 30 | N | N | 8 | exposed | deg1 | 2 | 3 |
| 31 | E | E | 13 | buried | deg1 | 2 | 3 |
| 32 | F | F | 9 | exposed | deg1 | 2 | 3 |
| 33 | R | R | 8 | exposed | deg1 | 2 | 3 |
| 34 | Y | Y | 10 | exposed | deg1 | 2 | 3 |
| 35 | F | F | 11 | buried | deg1 | 2 | 3 |
| 36 | Q | Q | 9 | exposed | deg1 | 2 | 3 |
| 37 | R | R | 8 | exposed | no | 2 | 5 |
| 38 | M | M | 10 | exposed | no | 2 | 5 |
| 39 | T | T | 12 | buried | no | 2 | 5 |
| 40 | T | T | 10 | exposed | no | 2 | 5 |
| 41 | T | T | 6 | exposed | no | 2 | 5 |
| 42 | S | S | 11 | buried | no | 2 | 5 |
| 43 | S | S | 5 | exposed | no | 2 | 5 |
| 44 | V | V | 6 | exposed | no | 2 | 5 |
| 45 | E | E | 4 | exposed | no | 2 | 5 |
| 46 | G | G | 6 | exposed | no | 2 | 5 |
| 47 | K | K | 10 | exposed | no | 2 | 5 |
| 48 | Q | Q | 11 | buried | no | 2 | 5 |
| 49 | N | N | 16 | buried | no | 4 | 5 |
| 50 | L | L | 13 | buried | no | 4 | 5 |
| 51 | V | V | 14 | buried | no | 4 | 5 |
| 52 | I | I | 15 | buried | no | 4 | 5 |
| 53 | M | M | 15 | buried | no | 4 | 5 |
| 54 | G | G | 13 | buried | no | 4 | 5 |
| 55 | K | K | 12 | buried | no | 4 | 5 |
| 56 | K | K | 8 | exposed | no | 4 | 5 |
| 57 | T | T | 12 | buried | no | 4 | 5 |
| 58 | W | W | 13 | buried | no | 4 | 5 |
| 59 | F | F | 8 | exposed | no | 4 | 5 |
| 60 | S | S | 8 | exposed | no | 4 | 5 |

| | | | | | | |
|---|---|---|---|---|---|---|
| 61 | I | I | 9 exposed | deg2 | 4 | 7 |
| 62 | P | P | 5 exposed | deg2 | 4 | 7 |
| 63 | E | E | 5 exposed | deg2 | 4 | 7 |
| 64 | K | K | 4 exposed | deg2 | 4 | 7 |
| 65 | N | N | 7 exposed | deg2 | 4 | 7 |
| 66 | R | R | 10 exposed | deg2 | 4 | 7 |
| 67 | P | P | 8 exposed | deg2 | 4 | 7 |
| 68 | L | L | 11 buried | deg2 | 4 | 7 |
| 69 | K | K | 6 exposed | deg2 | 4 | 7 |
| 70 | G | G | 6 exposed | deg2 | 4 | 7 |
| **71** | **R** | **R** | **13 buried** | **deg2** | **4** | **7** |
| **72** | **I** | **I** | **12 buried** | **deg2** | **4** | **7** |
| **73** | **N** | **N** | **14 buried** | **deg2** | **6** | **7** |
| **74** | **L** | **L** | **15 buried** | **deg2** | **6** | **7** |
| **75** | **V** | **V** | **15 buried** | **deg2** | **6** | **7** |
| **76** | **L** | **L** | **14 buried** | **deg2** | **6** | **7** |
| **77** | **S** | **S** | **9 exposed** | **deg2** | **6** | **7** |
| **78** | **R** | **R** | **3 exposed** | **deg2** | **6** | **7** |
| 79 | E | E | 4 exposed | deg2 | 6 | 7 |
| 80 | L | L | 9 exposed | deg2 | 6 | 7 |
| 81 | K | K | 2 exposed | deg2 | 6 | 7 |
| 82 | E | E | 5 exposed | deg2 | 6 | 7 |
| 83 | P | P | 10 exposed | deg2 | 6 | 7 |
| 84 | P | P | 11 buried | deg2 | 6 | 7 |
| 85 | Q | Q | 4 exposed | no | 6 | 9 |
| 86 | G | G | 8 exposed | no | 6 | 9 |
| 87 | A | A | 12 buried | no | 6 | 9 |
| 88 | H | H | 8 exposed | no | 6 | 9 |
| 89 | F | F | 11 buried | no | 6 | 9 |
| 90 | L | L | 12 buried | no | 6 | 9 |
| 91 | S | S | 11 buried | no | 6 | 9 |
| 92 | R | R | 7 exposed | no | 6 | 9 |
| 93 | S | S | 7 exposed | no | 6 | 9 |
| 94 | L | L | 13 buried | no | 6 | 9 |
| 95 | D | D | 6 exposed | no | 6 | 9 |
| 96 | D | D | 9 exposed | no | 6 | 9 |
| 97 | A | A | 13 buried | no | 8 | 9 |
| 98 | L | L | 11 buried | no | 8 | 9 |
| 99 | K | K | 8 exposed | no | 8 | 9 |
| 100 | L | L | 12 buried | no | 8 | 9 |
| 101 | T | T | 12 buried | no | 8 | 9 |
| 102 | E | E | 8 exposed | no | 8 | 9 |
| 103 | Q | Q | 8 exposed | no | 8 | 9 |
| 104 | P | P | 5 exposed | no | 8 | 9 |
| 105 | E | E | 6 exposed | no | 8 | 9 |
| 106 | L | L | 12 buried | no | 8 | 9 |
| 107 | A | A | 10 exposed | no | 8 | 9 |
| 108 | N | N | 6 exposed | no | 8 | 9 |
| 109 | K | K | 9 exposed | no | 8 | 11 |
| 110 | V | V | 15 buried | no | 8 | 11 |
| 111 | D | D | 10 exposed | no | 8 | 11 |
| 112 | M | M | 12 buried | no | 8 | 11 |
| 113 | V | V | 15 buried | no | 8 | 11 |
| 114 | W | W | 14 buried | no | 8 | 11 |
| 115 | I | I | 15 buried | no | 8 | 11 |
| 116 | V | V | 13 buried | no | 8 | 11 |
| 117 | G | G | 13 buried | no | 8 | 11 |
| 118 | G | G | 12 buried | no | 8 | 11 |
| 119 | S | S | 11 buried | no | 8 | 11 |
| 120 | S | S | 7 exposed | no | 8 | 11 |
| 121 | V | V | 15 buried | no | 10 | 11 |

| | | | | | | | |
|---|---|---|---|---|---|---|---|
| 122 | Y | Y | 14 | buried | no | 10 | 11 |
| 123 | K | K | 9 | exposed | no | 10 | 11 |
| 124 | E | E | 9 | exposed | no | 10 | 11 |
| 125 | A | A | 13 | buried | no | 10 | 11 |
| 126 | M | M | 10 | exposed | no | 10 | 11 |
| 127 | N | N | 5 | exposed | no | 10 | 11 |
| 128 | H | H | 9 | exposed | no | 10 | 11 |
| 129 | P | P | 3 | exposed | no | 10 | 11 |
| 130 | G | G | 8 | exposed | no | 10 | 11 |
| 131 | H | H | 7 | exposed | no | 10 | 11 |
| 132 | L | L | 13 | buried | no | 10 | 11 |
| 133 | K | K | 11 | buried | no | 10 | 13 |
| 134 | L | L | 15 | buried | no | 10 | 13 |
| 135 | F | F | 13 | buried | no | 10 | 13 |
| 136 | V | V | 12 | buried | no | 10 | 13 |
| 137 | T | T | 13 | buried | no | 10 | 13 |
| 138 | R | R | 10 | exposed | no | 10 | 13 |
| 139 | I | I | 15 | buried | no | 10 | 13 |
| 140 | M | M | 10 | exposed | no | 10 | 13 |
| 141 | Q | Q | 8 | exposed | no | 10 | 13 |
| 142 | D | D | 6 | exposed | no | 10 | 13 |
| 143 | F | F | 11 | buried | no | 10 | 13 |
| 144 | E | E | 6 | exposed | no | 10 | 13 |
| 145 | S | S | 12 | buried | no | 12 | 13 |
| 146 | D | D | 6 | exposed | no | 12 | 13 |
| 147 | T | T | 9 | exposed | no | 12 | 13 |
| 148 | F | F | 8 | exposed | no | 12 | 13 |
| 149 | F | F | 9 | exposed | no | 12 | 13 |
| 150 | P | P | 8 | exposed | no | 12 | 13 |
| 151 | E | E | 4 | exposed | no | 12 | 13 |
| 152 | I | I | 8 | exposed | no | 12 | 13 |
| 153 | D | D | 6 | exposed | no | 12 | 13 |
| 154 | L | L | 6 | exposed | no | 12 | 13 |
| 155 | E | E | 4 | exposed | no | 12 | 13 |
| 156 | K | K | 6 | exposed | no | 12 | 13 |
| 157 | Y | Y | 11 | buried | no | 12 | 15 |
| 158 | K | K | 6 | exposed | no | 12 | 15 |
| 159 | L | L | 8 | exposed | no | 12 | 15 |
| 160 | L | L | 9 | exposed | no | 12 | 15 |
| 161 | P | P | 3 | exposed | no | 12 | 15 |
| 162 | E | E | 6 | exposed | no | 12 | 15 |
| 163 | Y | Y | 6 | exposed | no | 12 | 15 |
| 164 | P | P | 4 | exposed | no | 12 | 15 |
| 165 | G | G | 4 | exposed | no | 12 | 15 |
| 166 | V | V | 9 | exposed | no | 12 | 15 |
| 167 | L | L | 6 | exposed | no | 12 | 15 |
| 168 | S | S | 6 | exposed | no | 12 | 15 |
| 169 | D | D | 5 | exposed | no | 14 | 15 |
| 170 | V | V | 7 | exposed | no | 14 | 15 |
| 171 | Q | Q | 11 | buried | no | 14 | 15 |
| 172 | E | E | 7 | exposed | no | 14 | 15 |
| 173 | E | E | 10 | exposed | no | 14 | 15 |
| 174 | K | K | 5 | exposed | no | 14 | 15 |
| 175 | G | G | 5 | exposed | no | 14 | 15 |
| 176 | I | I | 12 | buried | no | 14 | 15 |
| 177 | K | K | 10 | exposed | no | 14 | 15 |
| 178 | Y | Y | 15 | buried | no | 14 | 15 |
| 179 | K | K | 11 | buried | no | 14 | 15 |
| 180 | F | F | 13 | buried | no | 14 | 15 |
| 181 | E | E | 10 | exposed | no | 14 | 17 |
| 182 | V | V | 12 | buried | no | 14 | 17 |

| | | | | | | | |
|---|---|---|---|---|---|---|---|
| 183 | Y | Y | 12 | buried | no | 14 | 17 |
| 184 | E | E | 11 | buried | no | 14 | 17 |
| 185 | K | K | 12 | buried | no | 14 | 17 |
| 186 | N | N | 6 | exposed | no | 14 | 17 |
| 187 | D | D | 6 | exposed | no | 14 | 17 |
